# DSPE-PEG-CeO_2_ inhibits tumor cell proliferation by inducing oxidative stress and cell death

**DOI:** 10.1101/2023.07.26.550631

**Authors:** SiHao Qin, Xueyao Wang, Huan Li, YanFang Jiang

## Abstract

**SUMMARY:** CeO_2_ can consume intracellular reducing power, but its poor biocompatibility limits its application in tumor therapy. In this study, we designed and synthesized DSPE-PEG-CeO_2_, which is easy to enter cells, and showed that DSPE-PEG-CeO_2_ can induce the increase of ROS levels in tumor cells, thereby inducing tumor cell apoptosis and DNA damage. This suppresses the appearance and development of tumor cells. This provides a new idea for the application of inorganic nanomaterials.

## INTRODUCTION

It has been reported that nanocerium (CeO_2_), as a new type of nanozyme, can protect normal cells from radiation damage, and has the functions of superoxide dismutase, oxidase, catalase, and phosphatase^[1-6]^. Because of its enzyme-like activity, it plays an important role in environmental restoration, disease diagnosis, and biosensing. Although CeO_2_ has many advantages in applications, its easy aggregation and irregular distribution, especially in tumor killing.

The rapid proliferation of tumor cells usually faces three major challenges: biomacromolecule synthesis, energy supply, and oxidative stress^[7]^. Especially when oxidative stress accumulates in a large amount, it will destroy the integrity of intracellular biomacromolecules, such as lipids, enzyme activity, DNA structure, etc^[8]^. The most classic is the relationship between ROS and ferroptosis. A large number of studies have shown that the increase of intracellular ROS is the main factor of cell ferroptosis^[9]^. In addition, inducing ferroptosis in tumor cells has become an important means of tumor therapy^[10]^. Studies have shown that certain tumor cells are particularly sensitive to ferroptosis, such as fibrosarcoma, lung cancer, osteosarcoma, kidney cancer, and prostate cancer cells^[11-14]^.

Given the sensitivity of tumor cells to ROS and the potential role of CeO_2_ in regulating intracellular ROS, we hypothesized that CeO_2_ could be used to kill tumors. However, the easy aggregation and regular distribution of CeO_2_ limit its role in tumor therapy. However, as a widely used phospholipid-polymer conjugate, DSPE-PEG is expected to be an important adjuvant of CeO_2_ in tumor therapy. In this study, DSPE-PEG-CeO_2_ was designed and synthesized for the problem of easy aggregation of CeO_2_, and the material characterization test of DSPE-PEG-CeO_2_ was carried out. We then found that DSPE-PEG-CeO_2_ could induce an increase in ROS levels in tumor cells and induce cell death. However, it is worth noting that DSPE-PEG-CeO_2_ does not induce ferroptosis but inhibits cell proliferation by inducing apoptosis and DNA damage. At the same time, we confirmed in vivo experiments that DSPE-PEG-CeO_2_ can inhibit the growth of mouse tumors.

## MATERIALS AND METHODS

### Cell culture

HT1080 and HeLa were obtained from the Cell Bank of the Chinese Academy of Sciences (Shanghai, China). All mammalian cells were maintained in high glucose DMEM, sodium pyruvate, glutamine, penicillin, streptomycin, and 10% fetal bovine serum at 37°C and 5% CO_2_.

### Cell Viability Assay

For cell death experiments, cells were seeded in 96-well plates at 10,000 cells per well. 12 hours after sowing, they were treated with the indicated concentrations of DSPE-PEG-CeO_2_. Incubate with CCK8 for 1 hour and read at 450 nm.

### Measurement of cell death

Cells were seeded in 12-well culture dishes at a density of 1×10^5^ per well and cultured in DMEM for 12 hours. Add 1 μg/ml PI to the cell culture medium and incubate for 30 minutes. Labeled cells were trypsinized and resuspended in PBS and analyzed for cell death using PI-Stat coupled to flow cytometry.

### Measurement of ROS

ROS analysis by flow cytometry: Cells were seeded in 12-well dishes at a density of 1 × 10^5^/well and incubated in DMEM for 12 hours. Add 10 μM DCFH_2_-DA to the cell culture medium and incubate for 30 min. Subsequently, cells were washed twice with PBS to remove excess DCFH_2_-DA. Labeled cells were trypsinized and resuspended in PBS. ROS were analyzed using flow cytometry.

### Crystal violet staining

Cells were treated with the indicated concentrations of DSPE-PEG-CeO_2_ and cultured in DMEM medium. Colony formation was determined by crystal violet staining.

### Synthesis of DSPE-PEG-CeO_2_

Preparation of oil soluble cerium dioxide nanoparticles: 1.9 g of cerium acetate (Aladdin), 9.6 g of oleamide (Sigma), and 45 mL of xylene (Sigma) were placed in a 100 mL round bottom flask and stirred at room temperature for 24 hours to fully dissolve. Heat the oil bath to 90 ° C under argon protection; Quickly inject 3 mL of ddH_2_O with a syringe and react for 3 hours under reflux conditions. After the reaction is completed, slowly add the reaction solution under stirring conditions to 200 mL of acetone for precipitation at 4000rpm × Collect nanoparticles for 10 min. The concentrated particles were repeatedly washed with acetone (x5) to fully remove xylene. Finally, the particles are stored in chloroform for standby, and CeO_2_ nanoparticles with oleamine as ligand are prepared. DSPE-PEG coated oil soluble CeO_2_ to prepare water-soluble cerium dioxide nanoparticle liposomes: Disperse 30 mg of DSPE-PEG and 90 mg of oil soluble CeO_2_ in a 25 mL round bottom flask in 6 mL of chloroform, sonicate under ice bath conditions for 20 minutes, and then evaporate the solvent under reduced pressure distillation (60 ° C, about 30 minutes). The liposome CeO_2_ nanoparticles coated with DSPE-PEG were prepared by adding an appropriate amount of dd H_2_O (about 10 mL) and ultrasonic dispersion for 30 min. The crude product is placed in a dialysis bag (6000 Da) and dialyzed at room temperature with ddH_2_O for 72 hours. The dialysis solution is changed once every 12 hours. After completing dialysis purification, concentrate the particles using an ultrafiltration tube (3000 Da) and store them for future use.

### Particle characterization detection

Hydrated particle size, surface potential and stability tests in different solvents: DSPE-PEG-CeO_2_ was dispersed in 0.05 mg/mL ddH_2_O, and the hydrated particle size and zeta potential were measured using Zetasizer Nano. Measurement of UV-Vis absorption spectrum: DSPE-PEG-CeO_2_ was dispersed in ddH2O (0.05 mg/ml), and the absorption spectrum of the particles was measured with a UV-Vis spectrophotometer. Observation by transmission electron microscope: drop DSPE-PEG-CeO_2_ onto a copper grid to dry and observe the morphology of particles by transmission electron microscope.

### Xenograft Experiment

In xenografts bearing HeLa tumors, nude mice were injected subcutaneously with 5×10^6^ HeLa cells. Mice treated with PBS served as a control group and received intraperitoneal injections every 2 days from day 10 to day 25. All mice were euthanized, and tumors collected, weighed, embedded in paraffin, and further analyzed.

### Measurement of apoptosis

Flow Cytometry Analysis of Apoptosis: Cells were seeded in 12-well culture plates at a density of 1×10^5^/well and grown in DMEM for 12 hours. PI and annexin V were added to the cells, incubated for 30 min, and cell apoptosis was detected by flow cytometry.

### Statistical analysis

To ensure accuracy and reliability, all experiments were performed independently at least 3 times. Statistical analysis was performed using GraphPad Prism 6.0.1.

## RESULTS

### Preparation and characterization of DSPE-PEG-CeO_2_

To address the issue of cerium dioxide nanoparticles (CeO_2_) biocompatibility, we first prepared oil soluble CeO_2_, and then coated oil soluble CeO_2_ with DSPE-PEG to prepare water-soluble cerium dioxide nanoparticle liposomes (Fig. 1 A). Subsequently, we also observed the particle morphology of DSPE-PEG-CeO_2_ using a projection electron microscope, and the results showed that the particle morphology of DSPE-PEG-CeO_2_ was intact (Fig. 1 B).

**Fig 1.**
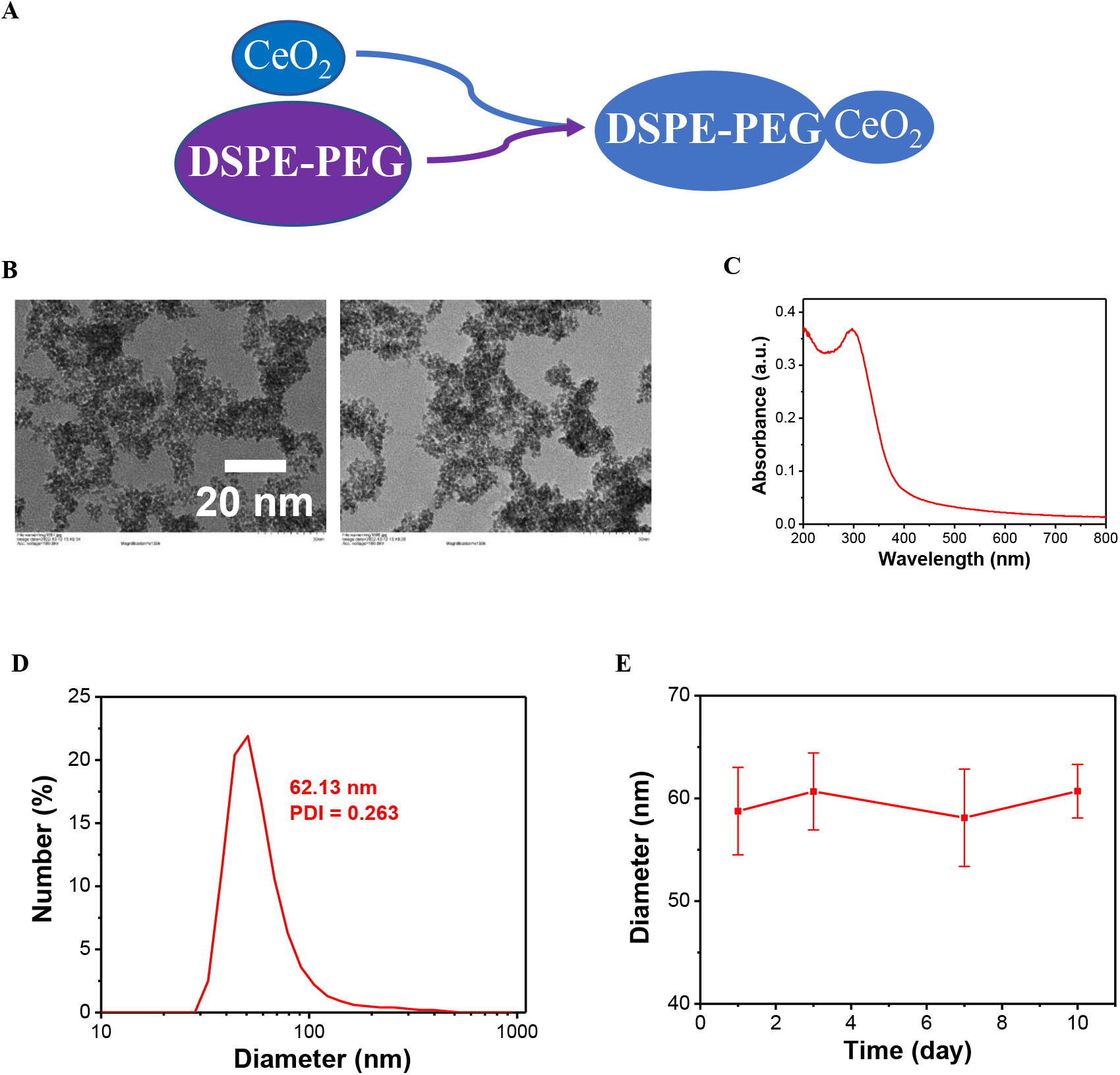
Preparation and characterization of DSPE-PEG-CeO_2_. (a) Design and synthesis diagram of DSPE-PEG-CeO_2_; (b) Drop the DSPE-PEG-CeO_2_ onto the copper mesh, dry them, and observe the particle morphology using a transmission electron microscope; (c) Disperse the DSPE-PEG-CeO_2_ in ddH2O (0.05 mg/ml) and measure the particle absorption spectrum using a PerkingElmer Lambda spectrophotometer; (d) (e) Disperse the DSPE-PEG-CeO_2_ in ddH_2_O (0.05 mg/ml) and measure the hydration particle size and Zeta potential using Zetasizer Nano (Malvin).

To address the biocompatibility of ceria nanoparticles (CeO_2_), we first prepared oil-soluble CeO_2_, and then coated oil-soluble CeO_2_ with DSPE-PEG to prepare water-soluble ceria nanoparticle liposomes (Fig. 1a). Subsequently, we also observed the particle morphology of DSPE-PEG-CeO_2_ with a projection electron microscope, and the results showed that the particle morphology of DSPE-PEG-CeO_2_was complete (Figure 1b). In addition, we dispersed DSPE-PEG-CeO_2_ in double distilled water and measured its hydrated particle size and zeta potential using Zetasizer Nano (Fig. 1c). The results show that the hydration particle size and zeta position of DSPE-PEG-CeO_2_ meet the design requirements. At the same time, we also obtained the absorption spectrum of DSPE-PEG-CeO_2_ with a Perking Elmer Lambda spectrophotometer, and the results showed that DSPE-PEG-CeO_2_ had an obvious absorption peak at 62.13 nm (Figure 1d and e)

### Cytotoxicity validation of DSPE-PEG-CeO_2_

In the above results, we synthesized DSPE-PEG-CeO_2_ and tested its material properties. The results show that the material meets the design requirements. Next, we will further test the cytotoxicity of DSPE-PEG-CeO_2_.We first treated HT1080 cells with different concentrations of DSPE-PEG-CeO_2_ for 42 hours. The results showed that 100 μg/mL DSPE-PEG-CeO_2_ could significantly inhibit tumor cell proliferation (Fig. 2a). Based on this, we investigated the treatment time of DSPE-PEG-CeO_2_, and the results showed that with the prolongation of DSPE-PEG-CeO_2_ treatment time, the proliferation of tumor cells was significantly inhibited (Figure 2b). In addition to examining the effect of DSPE-PEG-CeO_2_ on cell proliferation, we also examined the effect of DSPE-PEG-CeO_2_ treatment on the viability of CCK8 cells. The results showed that with the increase of A treatment concentration, the tumor cell viability gradually decreased (Fig. 2c). All the results indicated that DSPE-PEG-CeO_2_ cloud significantly inhibited tumor cell proliferation. However, our results indicated that DSPE-PEG-CeO_2_ could inhibit tumor cell proliferation by inducing tumor cell death.

**Fig 2.**
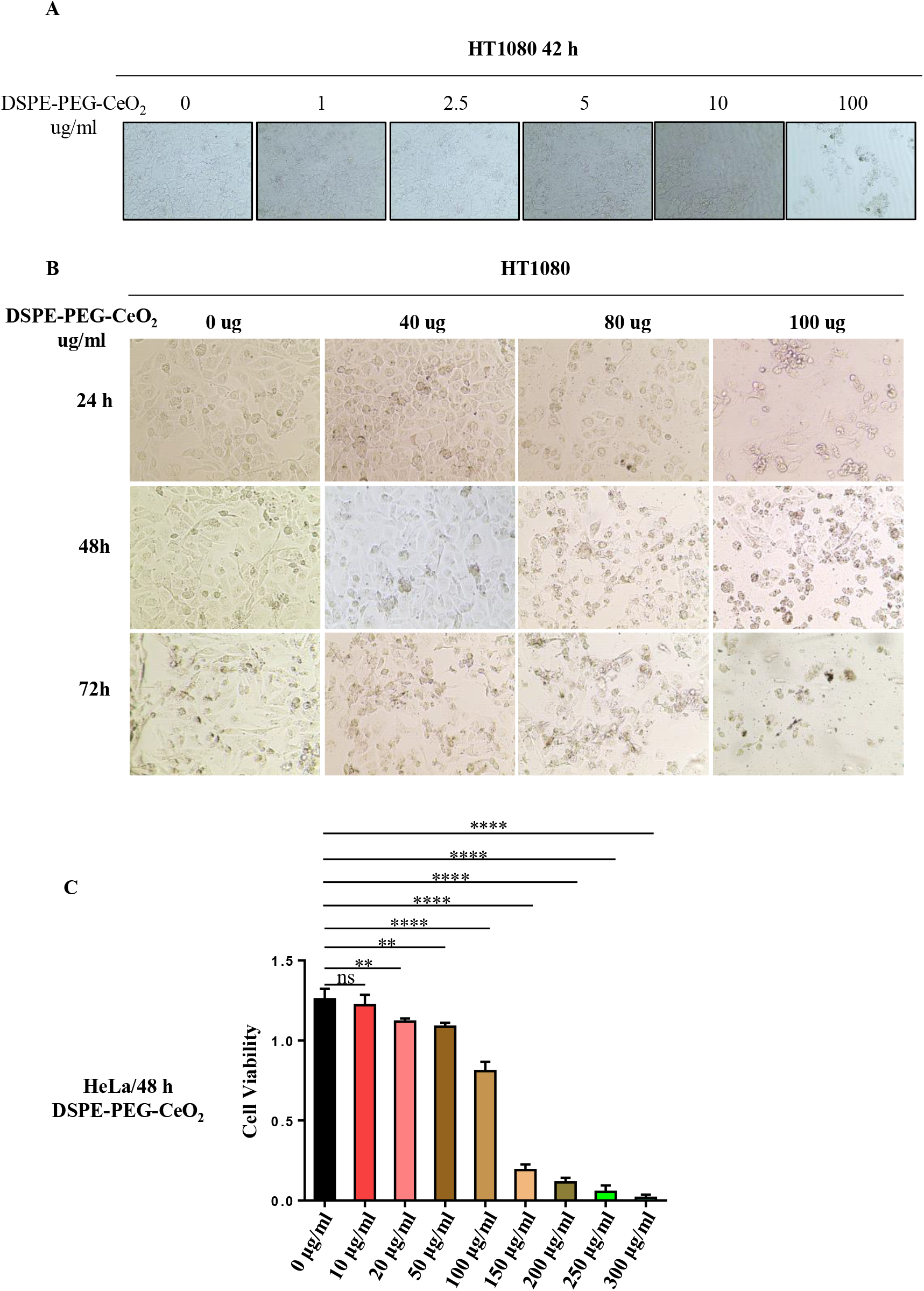
Cytotoxicity validation of DSPE-PEG-CeO_2_. (a) Treat HT1080 cells with different concentrations of DSPE-PEG-CeO_2_, and then examine the growth of multiple cells under microscope; (b) As shown, after treating HT1080 cells with DSPE-PEG-CeO_2_ for different times, the growth of the cells was examined under microscope; (c) After treating HT1080 cells with DSPE-PEG-CeO_2_, CCK detection of cell activity. ** <0.005, *** < 0.001, **** <0.0001.

### DSPE-PEG-CeO_2_ inhibits tumor cell proliferation by inducing cell death

To examine whether DSPE-PEG-CeO_2_ could induce tumor cell death, we first examined the effect of DSPE-PEG-CeO_2_ on HT1080 cell death by microscopy and crystal violet staining. The results showed that treatment with high concentrations of DSPE-PEG-CeO_2_ significantly induced tumor cell death (Fig. 3a and B). Meanwhile, we also used PI staining to detect cell death after DSPE-PEG-CeO_2_ treatment. The results showed that the cell death rate of HT1080 gradually increased with the concentration of DSPE-PEG-CeO_2_ treatment (Fig. 3c). Accordingly, we also counted the number of HeLa and HT1080 cells treated with DSPE-PEG-CeO_2_, and the results showed that DSPE-PEG-CeO_2_ significantly inhibited tumor cell proliferation (Figure 3d and e). Moreover, the proliferation of tumor cells is likely to achieved by inducing tumor cell death.

**Fig 3.**
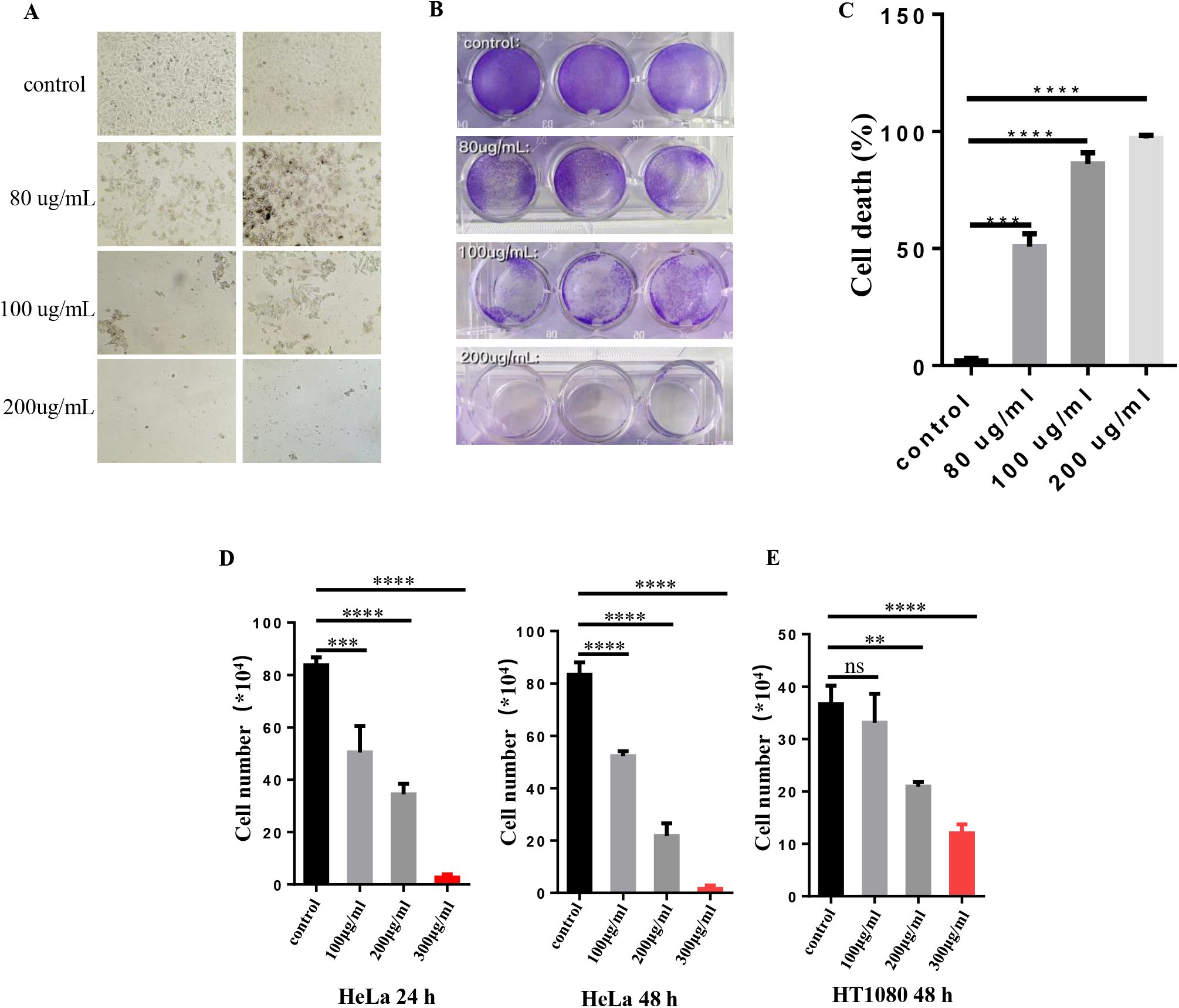
DSPE-PEG-CeO2 inhibits tumor cell proliferation by inducing cell death. (a) (b) After treating HT1080 cells with DSPE-PEG-CeO_2_, the cell growth was detected using a microscope and crystal violet staining; (c) After treating HT1080 cells with DSPE-PEG-CeO_2_, the cell death was detected using PI staining.; (d) (e) After treating HT1080 cells with DSPE-PEG-CeO_2_, count the number of cells. ** <0.005, *** <0.001, **** <0.0001.

### DSPE-PEG-CeO_2_ induces tumor cell death through non-ferroptosis

Considering that CeO_2_ has been reported to promote cellular oxidative stress by depleting intracellular NADPH, the above studies indicated that DSPE-PEG-CeO_2_ could induce tumor cell death. Therefore, we speculated that DSPE-PEG-CeO_2_ might induce cell death by increasing intracellular ROS levels. We treated HT1080 cells with DSPE-PEG-CeO_2_ to detect intracellular ROS levels. The results showed that DSPE-PEG-CeO_2_ significantly increased intracellular ROS levels (Fig. 4a). The accumulation of intracellular ROS has been reported to trigger lipid peroxidation, leading to cellular ferroptosis. We wanted to further confirm whether the increase of intracellular ROS induced by DSPE-PEG-CeO_2_ would induce ferroptosis. By microscopy and crystal violet staining, we found that DSPE-PEG-CeO_2_ could induce tumor cell death, but ferroptosis inhibitor could not inhibit DSPE-PEG-CeO_2_-induced tumor cell death (Fig. 4 b and c). Furthermore, we investigated the effect of DSPE-PEG-CeO_2_ and the ferroptosis inhibitor fer-1 on the viability of HeLa cells. The results showed that DSPE-PEG-CeO_2_ could limit the reduction of cell viability, but fer-1 did not alleviate this effect (Fig. 4 d). At the same time, the effect of DSPE-PEG-CeO_2_ on the ferroptosis of HeLa cells was proved. The results showed that intracellular lipid ROS did not increase after DSPE-PEG-CeO_2_ was applied to HeLa cells, and fer-1 could not inhibit DSPE-PEG-CeO2-induced cell death (Fig. 4e and f). In conclusion, DSPE-PEG-CeO_2_ can induce tumor cell death, but not tumor cell ferroptosis.

**Fig 4.**
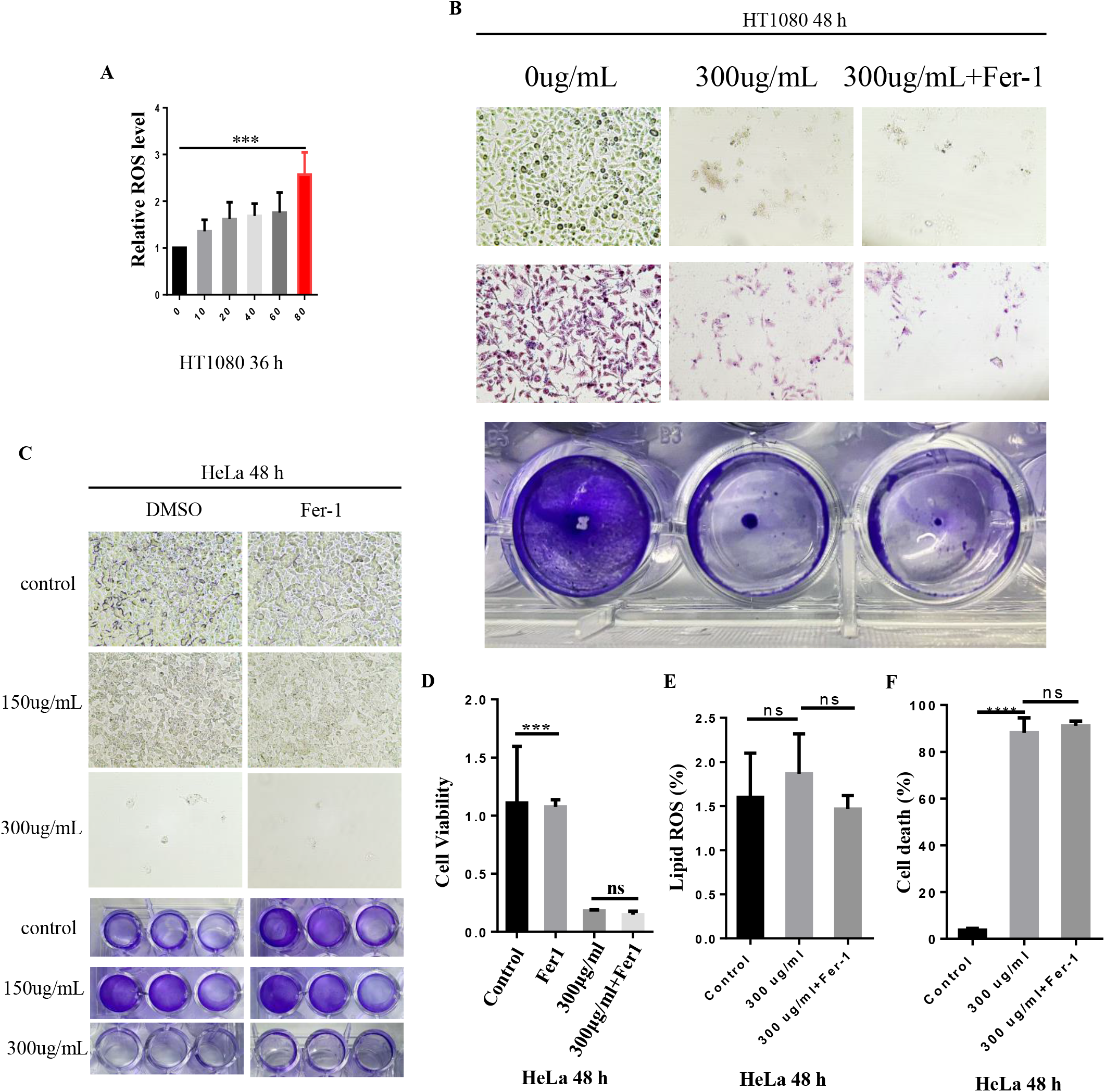
DSPE-PEG-CeO_2_ induces tumor cell death through non-ferroptosis. (a) After treating HT1080 cells with DSPE-PEG-CeO_2_, DCFH_2_-DA staining was used to detect intracellular ROS levels; (b) (c) After treating HT1080 cells with DSPE-PEG-CeO_2_ and Fer-1, Detection of cell growth using a microscope and crystal violet; (d) After treating HT1080 cells with DSPE-PEG-CeO_2_ and Fer-1, CCK8 detection of cell activity. (e) After treating HT1080 cells with DSPE-PEG-CeO2 and Fer-1, C11-BODIPY staining for Lipid ROS detection; (f) After treating HT1080 cells with DSPE-PEG-CeO2 and Fer-1, PI staining for cell death detection. ** < 0.005, *** <0.001, **** <0.0001.

### DSPE-PEG-CeO_2_ induces tumor cell death through apoptosis and DNA damage

In addition to triggering ferroptosis, ROS can also trigger apoptosis and DNA damage. Therefore, to further investigate the mechanism of DSPE-PEG-CeO_2_ induced cell death, we demonstrated the apoptosis of tumor cells after DSPE-PEG-CeO_2_ treatment. The results showed that DSPE-PEG-CeO_2_ could induce tumor cell apoptosis (Fig. 5a). At the same time, we also tested the effect of DSPE-PEG-CeO_2_ on DNA damage, and the results showed that DSPE-PEG-CeO_2_ could induce an increase in DNA damage index γ-H_2_A (Fig. 5b). Therefore, we also examined the effect of DSPE-PEG-CeO_2_ treatment on DNA replication efficiency and cell cycle. The results showed that DSPE-PEG-CeO_2_ could simultaneously inhibit DNA replication efficiency and cell cycle (Fig. 5c and d). These research results suggest that DSPE-PEG-CeO_2_ can induce tumor cell death by inducing apoptosis and DNA damage, and this effect is achieved by increasing intracellular ROS.

**Fig 5.**
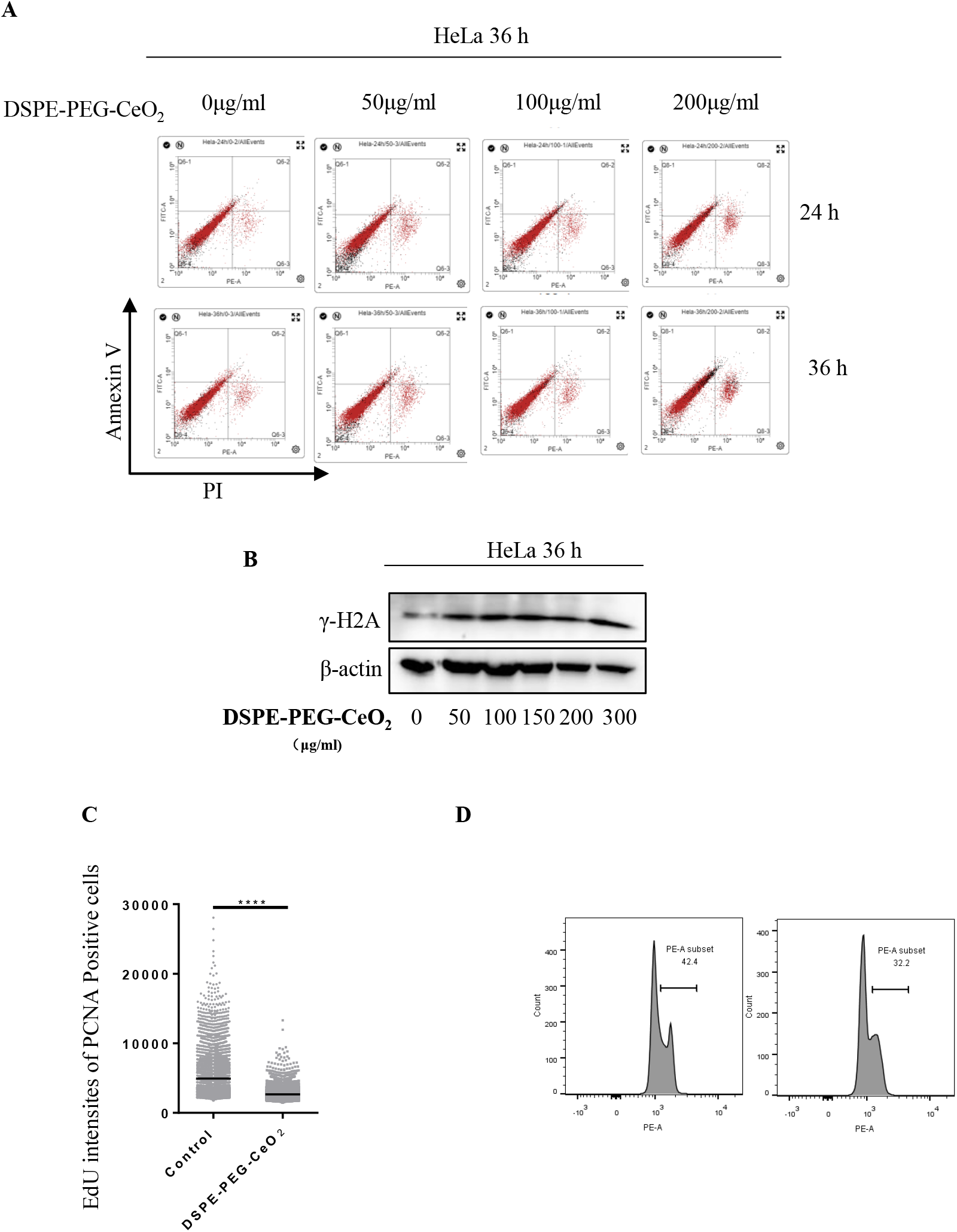
DSPE-PEG-CeO2 induces tumor cell death through apoptosis and DNA damage. (a) After treating HeLa cells with DSPE-PEG-CeO_2_, Detecting cell apoptosis; (b) After treating HeLa cells with DSPE-PEG-CeO_2_, WB detection of DNA damage; (c) After treating HeLa cells with DSPE-PEG-CeO_2_, test DNA replication efficiency. (d) After treating HeLa cells with DSPE-PEG-CeO2, detecting cell cycle. ** <0.005, *** <0.001, **** <0.0001.

### DSPE-PEG-CeO_2_ inhibit the proliferation of tumor cells in vivo

Through the above studies, we found that DSPE-PEG-CeO_2_ inhibited tumor cell proliferation by inducing an increase in ROS levels, thereby inducing tumor cell apoptosis and DNA damage. Then the antitumor effect of DSPE-PEG-CeO_2_ was tested on tumor-bearing nude mice. Nude mice were randomly divided into three groups (n=6) and were injected intravenously with PBS or DSPE-PEG-CeO_2_ (10 mg/kg). Mice treated with PBS served as a control group and received intraperitoneal injections every 2 days from day 10 to day 25. The results showed that compared with PBS administration, the DSPE-PEG-CeO_2_ group had the strongest antitumor effect (Fig. 6 a, b, c).

**Fig 6.**
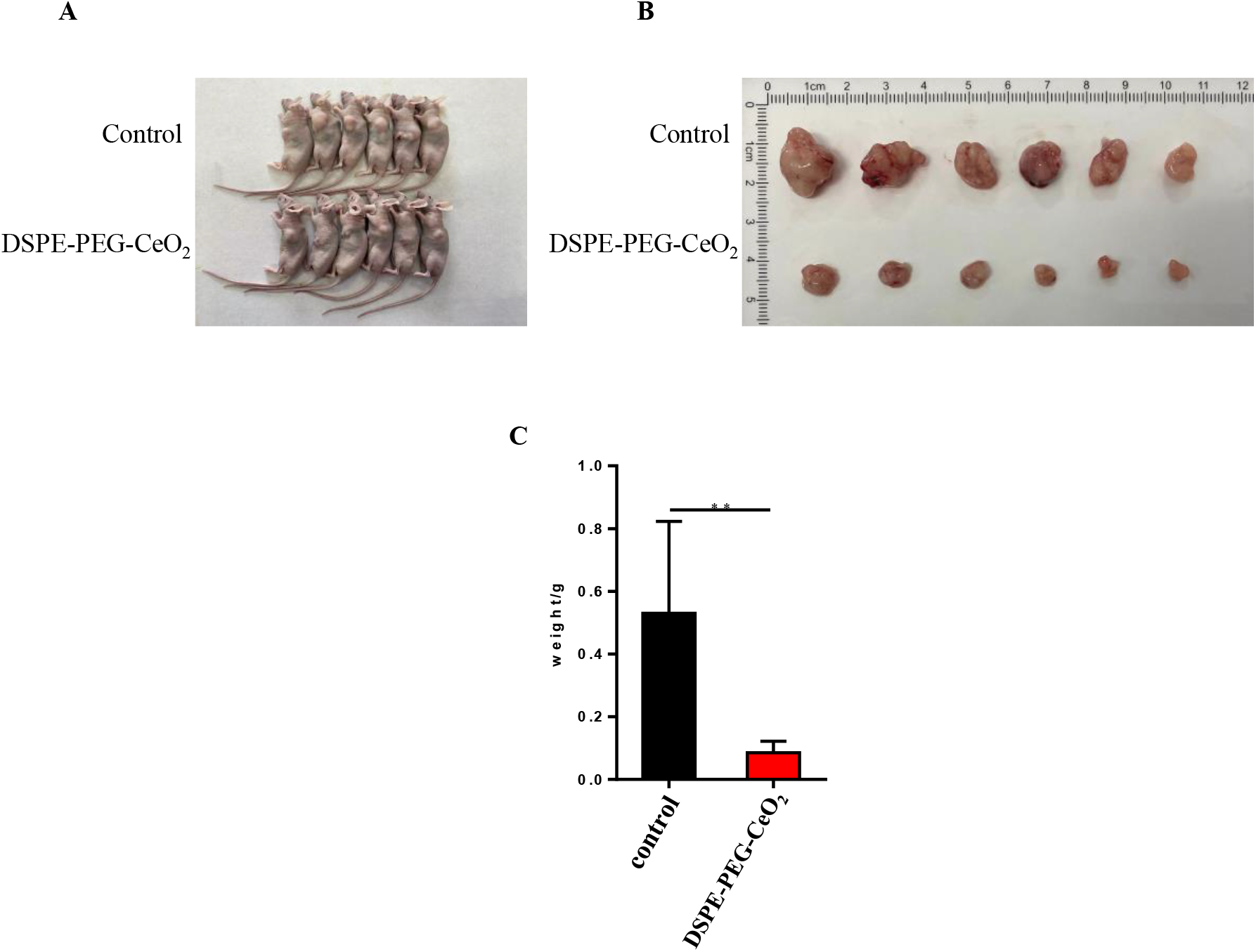
DSPE-PEG-CeO2 inhibit the proliferation of tumor cells in vivo. (a) Display tumor bearing mice and tumors; (b) Images of the collected tumors in different treatment groups on the 22nd day; (c) Weigh the collected tumors. ** <0.005.

## DISSUSSION

Inducing tumor cell death is still an important means of tumor treatment. The currently reported types of cell death are apoptosis, pyroptosis, ferroptosis, and copper death^[15, 16]^. Studies have shown that oxidative stress is an important means of inducing tumor cell death. For example, ROS can induce tumor cell ferroptosis by inducing lipid peroxidation^[17]^, and ROS can induce tumor cell apoptosis by inducing cytochrome *c* release^[18]^. The massive formation of intracellular ROS is mainly due to the disruption of the balance between intracellular oxidative power and reducing power. Therefore, the main way to induce the increase of ROS in tumor cells is to consume the reducing capacity in tumor cells. As an important nanozyme, one of its main properties is to consume intracellular NADPH, thereby increasing intracellular ROS. However, due to the poor biocompatibility of CeO_2_, its application in biology, especially in tumor killing, is limited^[19]^. Therefore, solving the problem of poor biocompatibility of nanozymes has become the key to the application of nanozymes in tumor therapy. And our work has just combined DSPE-PEG with CeO_2_, creatively solved the problem of poor biocompatibility of CeO_2_, and paved the way for its application in tumor therapy. In our work, we found that DSPE-PEG-CeO_2_ can induce tumor cell apoptosis and DNA damage by inducing the increase of ROS level to inhibit tumor cell proliferation. However, whether DSPE-PEG-CeO_2_ has other pathways to inhibit tumor cell proliferation and the limitations of its clinical application remain unclear.

## CONCLUSIONS

Overall, we designed and synthesized DSPE-PEG-CeO_2_to enhance the biocompatibility of CeO_2_. In vitro experiments showed that DSPE-PEG-CeO_2_ could inhibit tumor cell proliferation and induce cell death by increasing intracellular ROS levels and inducing DNA damage. We found that DSPE-PEG-CeO_2_ does not induce cell death by inducing ferroptosis in tumor cells. In vivo experiments, we found that DSPE-PEG-CeO2 can limit tumor proliferation. In summary, we applied DSPE-PEG-CeO_2_ to tumor therapy, which provided a successful argument for the application of nanozymes in anti-tumor therapy, and further expanded the application of nanozymes in biology.

## ACKNOWLEDGEMENTS

This research was funded by the National Natural Science Foundation of China (NOS 30972610 and 81273240), Jilin Province Science and Technology Agency (Nos JJKH20211210KJ, JJKH20211164KJ).

## AUTHOR CONTRIBUTIONS

J.Y., S.Q. conceived the project and designed the research; S.Q. carried out most of the experiments; X.W., H.L. carried out the mouse experiments; S.Q. analyzed the data; J.Y., S.Q. wrote the manuscript.

